# Temporal fine structure influences voicing confusions for consonant identification in multi-talker babble

**DOI:** 10.1101/2021.05.11.443678

**Authors:** Vibha Viswanathan, Barbara G. Shinn-Cunningham, Michael G. Heinz

## Abstract

To understand the mechanisms of speech perception in everyday listening environments, it is important to elucidate the relative contributions of different acoustic cues in transmitting phonetic content. Previous studies suggest that the envelope of speech in different frequency bands conveys most speech content, while the temporal fine structure (TFS) can aid in segregating target speech from background noise. However, the role of TFS in conveying phonetic content beyond what envelopes convey for intact speech in complex acoustic scenes is poorly understood. The present study addressed this question using online psychophysical experiments to measure the identification of consonants in multi-talker babble for intelligibility-matched intact and 64-channel envelope-vocoded stimuli. Consonant confusion patterns revealed that listeners had a greater tendency in the vocoded (versus intact) condition to be biased toward reporting that they heard an unvoiced consonant, despite envelope and place cues being largely preserved. This result was replicated when babble instances were varied across independent experiments, suggesting that TFS conveys voicing information beyond what is conveyed by envelopes for intact speech in babble. Given that multi-talker babble is a masker that is ubiquitous in everyday environments, this finding has implications for the design of assistive listening devices such as cochlear implants.

## 1 Introduction

Any acoustic signal can be decomposed into a slowly varying amplitude envelope, or temporal modulation, and a fast-varying temporal fine structure (TFS) (Hilbert, 1906). The cochlea decomposes sound input into a multi-channel representation organized by frequency, where each channel encodes the signal content in a relatively narrow band of frequencies around a given carrier frequency. The envelope and TFS information in each channel are then conveyed to the central nervous system through the ascending auditory pathway (Johnson, 1980; Joris and Yin, 1992). Elucidating the relative contributions of envelope and TFS cues to speech perception in everyday listening environments is important not just from a basic science perspective, but also for translation to clinical technologies such as cochlear implants.

Psychophysical studies suggest that speech content in quiet can be largely conveyed by envelopes (Shannon et al., 1995). Psychophysical (Bacon and Grantham, 1989; Stone and Moore, 2014), modeling (Dubbelboer and Houtgast, 2008; Relaño-Iborra et al., 2016), and electroencephalography (EEG) (Viswanathan et al., 2021) studies support the theory that in the presence of background noise, modulation masking of envelopes of target speech by distracting masker envelopes predicts speech intelligibility across diverse listening conditions. However, in addition to this contribution of envelopes to intelligibility, TFS may play a role, especially in noisy listening environments (Lorenzi et al., 2006; Hopkins and Moore, 2010).

Psychophysical studies suggest that cues conveyed by TFS (e.g., fundamental frequency (f0); Moore et al., 2006) can support perceptual scene segregation or unmasking (Darwin, 1997; Oxenham and Simonson, 2009). Moreover, EEG studies raised the possibility that the neural representation of the attended speech in a sound mixture is sensitive to the spectro-temporal details of the acoustic scene (Rimmele et al., 2015; Ding et al., 2014). By using high-resolution vocoding to alter TFS cues without introducing spurious envelopes, Viswanathan et al. (2021) showed that TFS cues *per se* can influence the coding of attended-speech envelopes in the brain and that this neural envelope coding in turn predicts intelligibility across a range of backgrounds and distortions. Despite the extensive prior literature on TFS and speech intelligibility, whether TFS can contribute to speech-in-noise perception beyond supporting masking release, i.e., whether TFS can directly convey phonetic content when envelopes are available, is poorly understood. As an analogy to help clarify this gap, consider the role of spatial cues. Spatial cues can provide masking release even though they do not carry any phonetic content. The analogous question here is whether TFS plays a similar role for speech perception in noise in that it only aids in unmasking or if TFS can also convey speech content when redundant intact envelope cues are available.

Previous behavioral studies that used TFS-vocoded speech (i.e., where the TFS or phase information in different frequency channels is retained but the envelope information is degraded; e.g., Sheft et al., 2008; Ardoint and Lorenzi, 2010) showed that TFS can convey certain phonetic features with relatively high levels of information reception by means other than envelope reconstruction (i.e., the recovery of degraded speech envelopes at the output of cochlear filters; Gilbert and Lorenzi, 2006; Heinz and Swaminathan, 2009). However, while these studies examined the role of TFS when envelope cues were degraded, they did not address the question of whether or not TFS cues are used for intact speech that has preserved envelope cues.

Another limitation of previous studies that investigated the role of TFS in conveying speech content is that they used masking conditions that were not ecologically realistic. While some used speech in quiet (Rosen, 1992; Sheft et al., 2008; Ardoint and Lorenzi, 2010), others presented speech in stationary noise (Gnansia et al., 2009; Swaminathan and Heinz, 2012). Ecologically relevant maskers such as multi-talker babble—a common source of interference in everyday cocktail-party listening—have not been utilized to study this problem. The spectro-temporal characteristics of multi-talker babble (envelope and TFS cues) are similar to what may be encountered in realistic scenarios and a better match to competing speech (albeit without semantic and linguistic content). Thus, multi-talker babble is an important masker to use when studying the role of TFS in speech understanding.

The present study addressed these gaps using online envelope-vocoding experiments designed to probe directly the role of TFS in conveying consonant information beyond what envelopes convey for intact speech (i.e., with redundant envelope cues) in realistic masking environments. Multi-talker babble was used as an ecologically relevant masker. Consonant confusion patterns (Miller and Nicely, 1955) were analyzed, grouping consonants into categories based upon the features of voicing, place of articulation (POA), and manner of articulation (MOA). Confusion patterns were compared between intact and 64-channel envelope-vocoded stimuli for consonants presented in multi-talker babble and separately in quiet (as a control). 64-channel envelope vocoding largely preserves cochlear-level envelopes (Viswanathan et al., 2021), allowing us to study the role of the original TFS in conveying speech content beyond what is conveyed by the intact envelopes. Since TFS plays a role in masking release, vocoding at the same signal-to-noise ratio (SNR) as for intact stimuli produces considerably lower intelligibility. Here, this intelligibility drop was mitigated by using a higher SNR for vocoded stimuli so that overall intelligibility was matched for intact and vocoded conditions. By matching intelligibility in this manner, differences in confusion patterns across conditions could be attributed to changes in consonant categorization and category errors rather than differences in overall error counts. Moreover, equalizing intelligibility maximized the statistical power for detecting differences in the pattern of confusions. Finally, given that consonants are transient sounds, whether or not effects were robust to changes in the local statistics of the masker was also examined by testing whether results were replicated when the specific instances (i.e., realizations) of multi-talker babble varied across experiments.

The current study tested the hypothesis that TFS does not convey speech content beyond what is conveyed by envelopes for intact speech (i.e., the classic view that envelopes convey speech content and that TFS conveys other attributes like pitch and aids source segregation). If this is the case, then once intelligibility is matched across conditions, confusion patterns should be the same for intact and envelope-vocoded stimuli corresponding to speech in (i) babble and (ii) quiet. The experiments used to test this hypothesis, the results, and their implications are described below.

## 2 Materials and Methods

### 2.1 Stimulus generation

Twenty consonants from the STeVI corpus (Sensimetrics Corp., Malden, MA) were used. The consonants were /b/, /t∫/, /d/, /ð/, /f/, /g/, /d3/, /k/, /l/, /m/, /n/, /p/, /r/, /s/, /∫/, /t/, /θ/, /v/, /z/, and /3/. The consonants were presented in consonant-vowel (CV) context, where the vowel was always /a/. Each consonant was spoken by two female and two male talkers (to reflect real-life talker variability). The CV utterances were embedded in the carrier phrase: “You will mark /CV/ please” (i.e., in natural running speech). Stimuli were created for five experimental conditions:

1. **Speech in babble (SiB):** Speech was added to four-talker babble at -8 dB SNR. The long-term spectrum of the target speech (including the carrier phrase) was adjusted to match the average (across instances) long-term spectrum of the four-talker babble (by applying a filter with a transfer function equal to the ratio of the two spectra). To create each SiB stimulus, a babble sample was randomly selected from a list comprising 72 different four-talker babble maskers obtained from the QuickSIN corpus (Killion et al., 2004).
2. **Vocoded speech in babble (vocoded SiB):** SiB at 0 dB SNR was subjected to 64-channel envelope vocoding. A randomly selected babble sample was used for each vocoded SiB stimulus, similar to what was done for intact SiB. The vocoding process retained the cochlear-level envelopes but replaced the stimulus fine structure with a noise carrier, in accordance with the procedure described in Qin and Oxenham (2003). The 64 frequency channels were contiguous with their center frequencies equally spaced on an equivalent rectangular bandwidth (ERB)-number scale (Glasberg and Moore, 1990) between 80 and 6000 Hz. This resulted in roughly two channels per ERB, which ensured that for any given channel, there was one additional channel on each side within 1 ERB. This helps to mitigate spurious envelope recovery on the slopes of cochlear filters, which in turn allows for TFS effects to be better isolated (Viswanathan et al., 2021). The envelope in each channel was extracted using a sixth-order Butterworth band-pass filter followed by half-wave rectification and low-pass filtering using a second-order Butterworth filter with a cutoff frequency of 300 Hz or half of the channel bandwidth, whichever was lower. The envelope in each channel was then used to modulate a random Gaussian white noise carrier; the result was band-pass filtered within the channel bandwidth and scaled to match the level of the original signal.
3. **Speech in quiet (SiQuiet):** Speech in quiet was used as a control condition.
4. **Vocoded speech in quiet (vocoded SiQuiet):** SiQuiet subjected to 64-channel envelope vocoding (using the same procedure as for vocoded SiB) was used to examine whether TFS conveys speech content beyond what envelopes convey for intact speech in quiet.
5. **Speech in speech-shaped stationary noise (SiSSN):** Speech was added to stationary Gaussian noise at -8 dB SNR. Similar to what was done for SiB, the long-term spectra of the target speech (including the carrier phrase) and that of the stationary noise were adjusted to match the average (across instances) long-term spectrum of the four-talker babble. A different realization of stationary noise was used for each SiSSN stimulus. The SiSSN condition was used for online data quality checking, given that lab-based confusion data were available for this condition (Phatak and Allen, 2007).

Prior to the main consonant identification study, a behavioral pilot study (with three subjects who did not participate in the actual online experiments) was used to determine appropriate SNRs for the different experimental conditions. The SNRs for the intact and vocoded SiB conditions were chosen to give intelligibility of roughly 60%, so that a sufficient number of confusions would be obtained for data analysis.

To verify that the vocoding procedure did not significantly change envelopes at the cochlear level, the envelopes at the output of 128 filters were extracted (using a similar procedure as in the actual vocoding process) both before and after vocoding for SiQuiet and SiB at 0 dB SNR and for each of the different consonants and talkers. The use of 128 filters allowed us to compare envelopes for both on-band filters (i.e., filters whose center frequencies matched those of the subbands of the vocoder) and off-band filters (i.e., filters whose center frequencies were halfway between adjacent vocoder subbands on the ERB-number scale). The average correlation coefficient between envelopes before and after vocoding (across the different stimuli and cochlear filters and after adjusting for any vocoder group delays) was about 0.9 (Fig. 1). This suggests that the 64-channel envelope-vocoding procedure left the within-band cochlear-level envelopes largely intact. Thus, although intrinsic envelope fluctuations conveyed by the noise carrier used in vocoding may mask crucial speech-envelope cues in some cases (Kates, 2011), this issue is mitigated by using high-resolution vocoding as was done in the current study. This high-resolution vocoding allowed us to unambiguously attribute vocoding effects to TFS cues rather than any spurious envelopes (not present in the original stimuli) that can be introduced within individual frequency bands during cochlear filtering of the noise carrier used in vocoding when low-resolution vocoding is performed (Gilbert and Lorenzi, 2006; Swaminathan and Heinz, 2012; Viswanathan et al., 2021).

**Figure 1.**
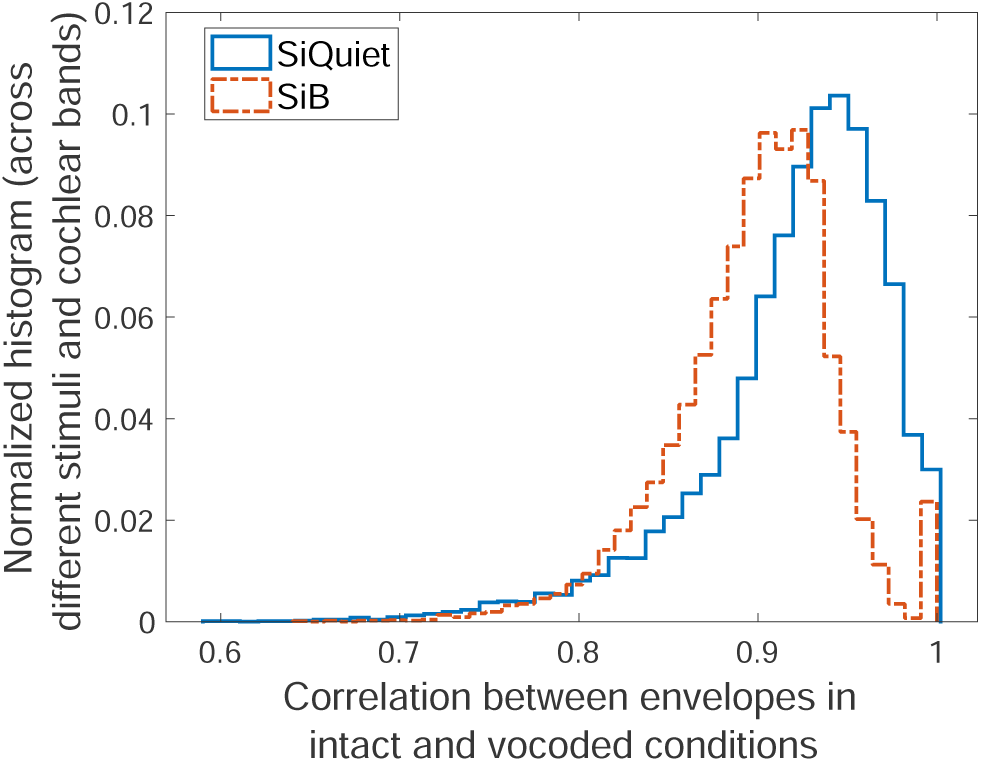
64-channel envelope vocoding largely preserves the envelopes within individual cochlear bands. Shown are the normalized histogram of the group-delay-adjusted correlation between the envelope for intact speech in quiet (SiQuiet) and 64-channel vocoded SiQuiet (i), and that for intact speech in babble (SiB) and 64-channel vocoded SiB (ii). The histograms are across the different consonants and talkers as well as across 128 different cochlear bands equally spaced on an ERB-number scale from 80 to 6000 Hz. The average correlation between envelopes before and after vocoding was about 0.9.

The stimulus used for online volume adjustment was separately generated and consisted of running speech mixed with four-talker babble. The speech and babble samples were both obtained from the QuickSIN corpus (Killion et al., 2004); these were repeated over time to obtain a total stimulus duration of ∼20 s (to give subjects adequate time to adjust their computer volume with the instructions described in Section 2.3). The volume-adjustment stimulus was designed to have a root mean square (RMS) value that corresponded to 75% of the dB difference between the softest and loudest stimuli in the study. This ensured that once subjects had adjusted their computer volume, the stimuli used in the main consonant identification tasks were never too loud for subjects, even at adverse SNRs.

### 2.2 Participants

Data were collected online from anonymous subjects recruited using Prolific.co. The subject pool was restricted using a screening method developed by Mok et al. (2021). The screening method contained three parts: (i) a core survey that was used to restrict subjects based on age to 18–55 years (to exclude significant age-related hearing loss), whether or not they were US/Canada residents, US/Canada born, and native speakers of North American English (because North American speech stimuli were used), history of hearing and neurological diagnoses if any, and whether or not they had persistent tinnitus; (ii) headphone/earphone checks; and (iii) a speech-in-babble-based hearing screening. Subjects who passed the screening were invited to participate in the consonant identification study, and when they returned, headphone/earphone checks were performed again. All subjects had completed at least 40 previous studies on Prolific and had *>* 90% of them approved (Prolific allows researchers to reject participant submissions if there is clear evidence of non-compliance with instructions or poor attention). These procedures were validated in previous work, where they were shown to successfully select participants with near-normal hearing status, attentive engagement, and stereo headphone use (Mok et al., 2021). Subjects provided informed consent in accordance with remote testing protocols approved by the Purdue University Institutional Review Board (IRB).

### 2.3 Experimental design

Three nearly identical consonant-identification experiments were conducted to assess the replicability of any main effect of TFS. The experiments were designed with the goal of contrasting intact and vocoded conditions (i.e., stimuli with original and disrupted TFS) while roving the levels of all other experimental variables (i.e., consonants, talkers, conditions, and masker instances). Thus, each experiment presented, in random order, one stimulus repetition for each of the 20 consonants across all four talkers and all five experimental conditions. Within a given experiment, in creating each intact or vocoded SiB stimulus, babble instances (i.e., realizations) were randomly chosen from a list comprising 72 different four-talker babble maskers (see Section 2.1); thus, the babble instances that were used for a particular consonant and talker were not the same between the intact and vocoded SiB conditions. To test whether the main effects of TFS generalized when the babble instances used were varied across experiments, a different random pairing of masker instances was used across consonants, talkers, and conditions in experiment 2 compared to experiment 1. Experiment 3 used, as a sanity check while testing replication of effects, the same stimuli as experiment 2. Thus, the only difference in the stimuli between the experiments was in the particular instance of babble that was paired with a particular consonant, talker, and SiB condition (intact and vocoded). As observed by Zaar and Dau (2015), when effects are instance-specific, different realizations of the same masker random process can contribute significantly larger variability to consonant identification measurements than across-listener variability. Thus, our study design of varying babble instances across the three experiments helped to disambiguate any effects of vocoding from masker-instance effects.

Twenty-five subjects per talker were used (subject overlap between talkers was not controlled) in each of the three experiments. With four talkers, this yielded 100 subject-talker pairs, or samples, per experiment. Separate studies were posted on Prolific.co for the different talkers; thus, when a subject performed a particular study, they would be presented with the speech stimuli for one specific talker consistently over all trials. There was no overlap between experiments in the particular set of 100 samples that were used, i.e., samples were independent across experiments. Within each experiment, talker, and condition, all subjects performed the task with the same stimuli. Moreover, all condition effect contrasts were computed on a within-subject basis and averaged across subjects.

Subjects performed the tasks using their personal computers and headphones/earphones. Our online infrastructure included checks to prevent the use of mobile devices. Each of the three experiments had three parts: (i) Headphone/earphone checks, (ii) Demonstration (“demo”), and (iii) test (which was the main stage of the experiment). Each of these three parts had a volume-adjustment task at the beginning. In this task, subjects were asked to make sure that they were in a quiet room and wearing wired (not wireless) headphones or earphones. They were instructed not to use desktop/laptop speakers. They were then asked to set their computer volume to 10%–20% of the full volume, following which they were played a speech-in-babble stimulus and asked to adjust their volume up to a comfortable but not too loud level. Once subjects had adjusted their computer volume, they were instructed not to adjust the volume during the experiment, as that could lead to sounds being too loud or soft.

The paradigm of Mok et al. (2021) was used for headphone/earphone checks. In this paradigm, subjects first performed the task described by Woods et al. (2017). While the Woods et al. (2017) task can distinguish between listening with a pair of free-field speakers versus using stereo headphones/earphones, it cannot detect the use of a single free-field speaker or a mono headphone/earphone. Thus, the Woods et al. (2017) task was supplemented with a second task where the target cues were purely binaural in nature, thereby allowing us to test if headphones/earphones were used in both ears. The second task was a three-interval three-alternative forced-choice task where the target interval contained white noise with interaural correlation fluctuating at 20 Hz, while the dummy intervals contained white noise with a constant interaural correlation. Subjects were asked to detect the interval with the most flutter or fluctuation. Only those subjects who scored greater than 65% in each of these two tasks were allowed to proceed to the next (demo) stage of the experiment. This two-task paradigm to verify stereo headphone/earphone use was validated in Mok et al. (2021).

In the demo stage, subjects performed a short training task designed to familiarize them with how each consonant sounds and with the consonant-identification paradigm. Subjects were instructed that in each trial they would hear a voice say “You will mark *something* please.” They were told that at the end of the trial, they would be given a set of options for *something* and that they would have to click on the corresponding option. Consonants were first presented in quiet and in sequential order starting with /b/ and ending with /3/. This order was matched in the consonant options shown on the screen at the end of each trial. After the stimulus ended in each trial, subjects were asked to click on the consonant they heard. After subjects had heard all consonants sequentially in quiet, they were tasked with identifying consonants presented in random order and spanning the same set of listening conditions as the test stage. Subjects were instructed to ignore any background noise and only listen to the particular voice saying “You will mark *something* please.” Only subjects who scored ≥ 85% in the demo’s SiQuiet control condition were selected for the test stage so as to ensure that all subjects understood and were able to perform the task.

In the test stage, subjects were given similar instructions as in the demo but told to expect trials with background noise from the beginning (rather than midway through the task as in the demo). In both demo and test, the background noise (babble or stationary noise), when present, started 1 s before the target speech and continued for the entire duration of the trial. In both demo and test, to promote engagement with the task, subjects received feedback after every trial as to whether or not their response was correct. Subjects were not told what consonant was presented, to avoid over-training to the acoustics of how each consonant sounded across the different conditions, except for the first sub-part of the demo, where subjects heard all consonants in quiet in sequential order.

### 2.4 Data preprocessing

Only samples (i.e., subject-talker pairs) with intelligibility scores ≥ 85% for the SiQuiet control condition in the test stage were included in results reported here. All conditions for the remaining samples were excluded from further analyses as a data quality control measure.

### 2.5 Quantifying confusion matrices

The 20 English consonants used in this study were assigned the phonetic features described in Table 1. The identification data collected in the test stage of each experiment were used to construct consonant confusion matrices (pooled over samples) separately for each condition. Overall intelligibility was normalized to 60% for intact and vocoded SiB and to 90% for intact and vocoded SiQuiet by scaling the confusion matrices such that the sum of the diagonal entries was the desired intelligibility. Matching intelligibility in this manner allowed for differences in confusion patterns across conditions to be attributed to changes in consonant categorization and category errors rather than differences in overall error counts (due to one condition being inherently easier at a particular SNR). Furthermore, equalizing intelligibility maximizes the statistical power for detecting differences in the pattern of confusions. The resulting confusion matrices (Fig. 9) were used to construct voicing, POA, and MOA confusion matrices by pooling over all consonants. To test our hypothesis that voicing, POA, and MOA confusion patterns would be the same for intact and envelope-vocoded speech in babble (after matching intelligibility), the difference between intelligibility-matched intact and vocoded SiB confusion matrices was computed. Confusion-matrix differences were then compared with appropriate null distributions of zero differences (see Section 2.6) to extract statistically significant differences (shown in Figs. 6, 7, and 8). A similar procedure was used to test whether TFS conveys phonetic content beyond what is conveyed by envelopes for intact speech in quiet, but by pooling data across all three experiments when constructing confusion matrices for intact and vocoded SiQuiet (versus examining effects separately for each experiment, as was done for intact and vocoded SiB). This data pooling across experiments was performed to improve statistical power because of the relatively high overall intelligibility in quiet.

**Table 1.**
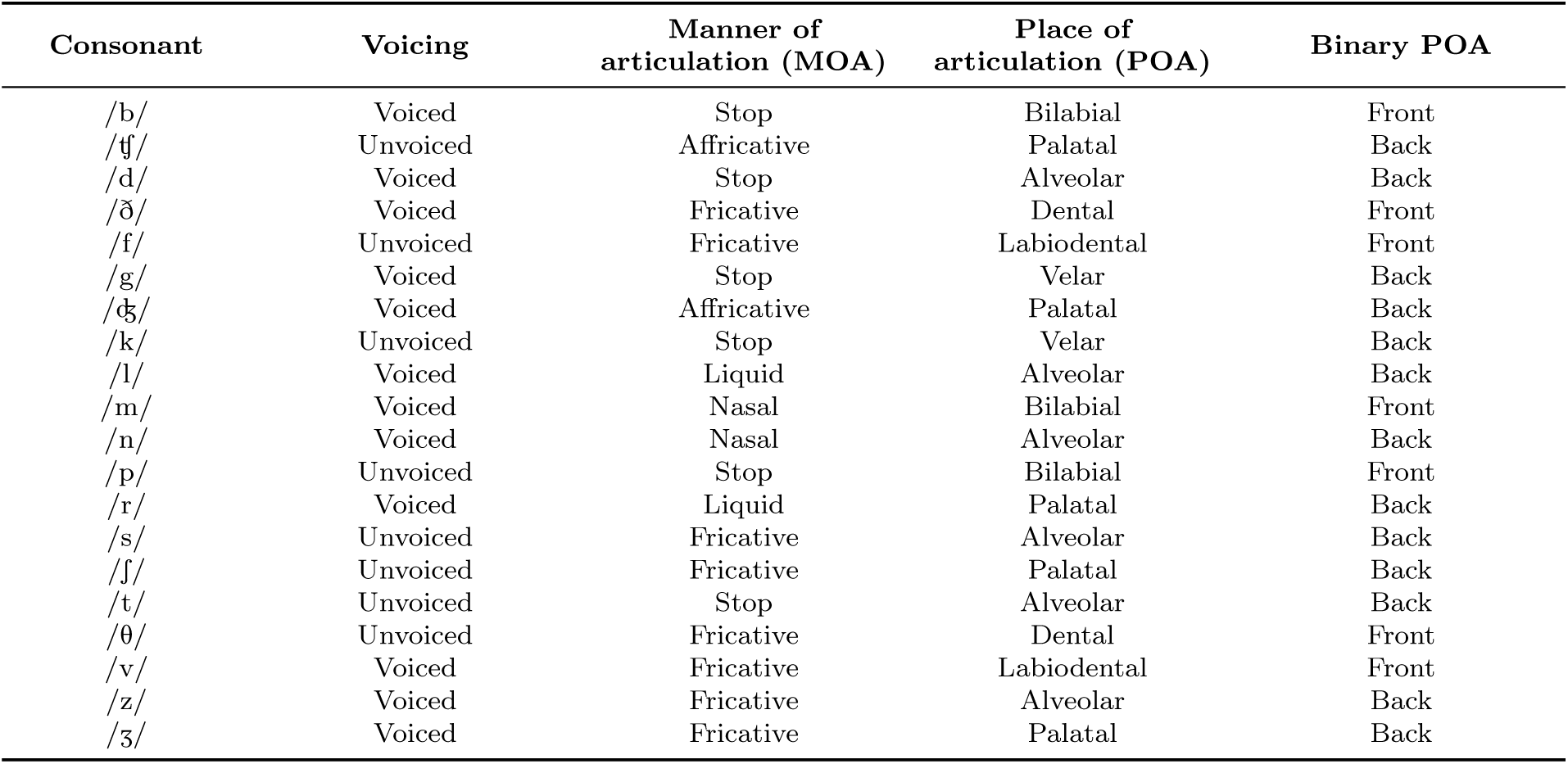
Phonetic features of the 20 English consonants used in this study.

### 2.6 Statistical analysis

To examine the role of TFS in conveying speech content, the difference in the voicing, POA, and MOA confusion matrices between intact and vocoded conditions was computed, separately for speech in babble and speech in quiet. Permutation testing (Nichols and Holmes, 2002) with multiple-comparison correction at 5% false-discovery rate (FDR; Benjamini and Hochberg, 1995) was used to extract significant differences in the confusion patterns. The null distributions for permutation testing were obtained using a non-parametric shuffling procedure, which ensured that the data used in the computation of the null distributions had the same statistical properties as the measured confusion data. Separate null distributions were generated for speech in babble and speech in quiet, and for the different phonetic categories. Each realization from each null distribution was obtained by following the same computations used to obtain the actual “intact - vocoded” confusion matrices but with random shuffling of intact versus vocoded condition labels corresponding to the measurements. This procedure was repeated with 10,000 distinct randomizations for each null distribution.

To quantify the degree to which statistically significant “intact - vocoded” confusion differences were replicated across the three experiments, simple Pearson correlation was used and the p-value for the correlation was derived using Fisher’s approximation (Fisher, 1921). Although the entries of each difference matrix are not strictly independent (which can cause p-values to be underestimated), this p-value approximation was considered adequate given that the individual p-value estimates were not near conventional significance criteria (i.e., were orders of magnitude above or below 0.05).

### 2.7 Signal-detection theoretic analysis

A signal-detection theoretic analysis (Green and Swets, 1966) was used to calculate the bias, i.e., the shift in the classification boundary, in the average subject’s percept of voicing for target speech in babble relative to an unbiased ideal observer (i.e., a classifier that optimally uses the acoustics to arrive at a speech-category decision) (see Fig. 2). The extent to which this bias was altered by vocoding was then quantified. This analysis was motivated by the finding that vocoding had a significant and replicable effect on voicing confusions for speech in babble across the three experiments in our study.

**Figure 2.**
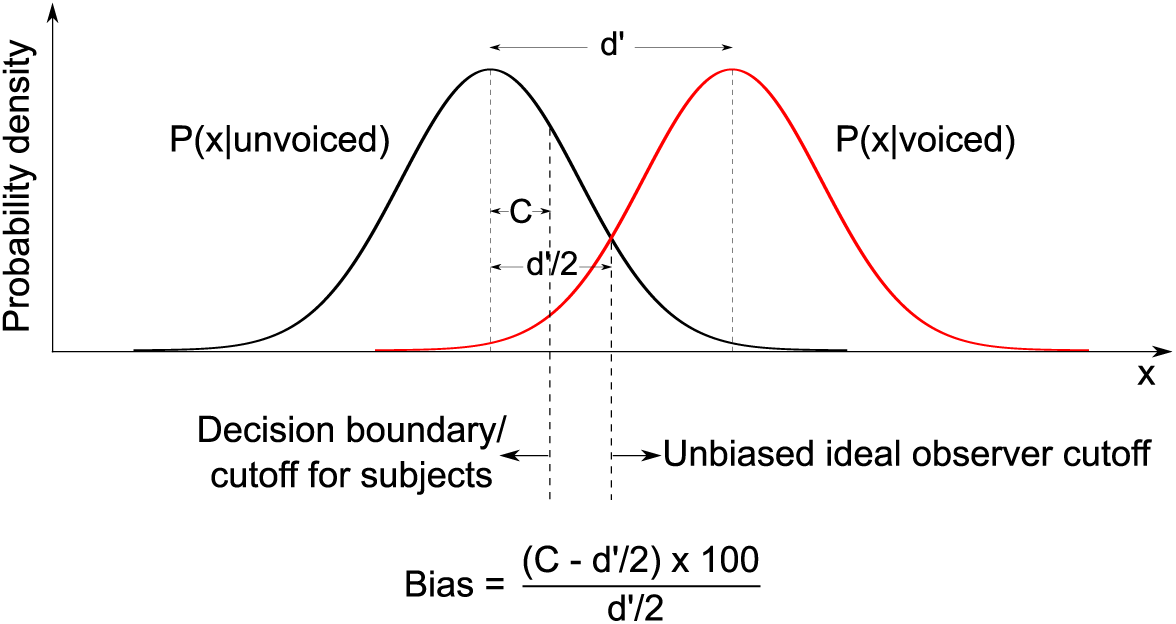
Illustration of a decision-theoretic quantification of speech categorization bias. *x* denotes the internal decision variable. Bias is quantified as the percent shift in the average listener’s cutoff (or decision boundary) relative to an unbiased ideal observer’s cutoff. The cutoff values for the average listener and the ideal observer were estimated from the false-alarm and hit rates in the data.

Let us define the null and alternative hypotheses for the voicing categorization performed by listeners. Let ℋ0 be the null hypothesis that an unvoiced consonant was presented, and let ℋ1 be the alternative hypothesis that a voiced consonant was presented. Let *FA* be the probability of a false alarm, and let *HR* be the hit rate. The *FA* and *HR* values for each experiment and condition were obtained from the voicing confusion matrix (pooled over samples and consonants) corresponding to that experiment and condition.

The cutoff *C* (or decision boundary) for the average subject’s perceptual decision on whether or not to reject ℋ0, *d*′, and listener bias *B* (expressed as a percentage relative to an unbiased ideal observer’s cutoff) were calculated separately for each experiment and condition (intact versus vocoded SiB) as:

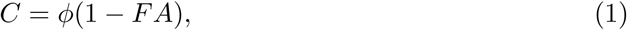

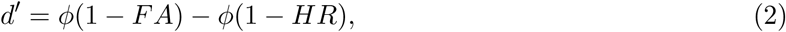

and

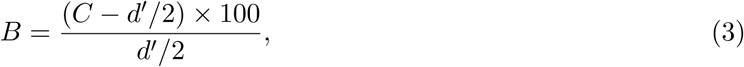

where *ϕ* is the inverse of the standard normal cumulative distribution.

The change in the listener bias between the intact and vocoded SiB conditions was derived as:

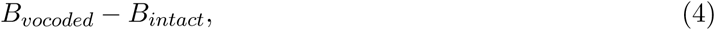

where *B*_*vocoded*_ and *B*_*intact*_ are the biases in the vocoded and intact SiB conditions, respectively.

### 2.8 Software accessibility

Subjects were directed from Prolific to the SNAPlab online psychoacoustics infrastructure (Bharadwaj, 2021; Mok et al., 2021) to perform the study. Offline data analyses were performed using custom software in PYTHON (Python Software Foundation, Wilmington, DE) and MATLAB (The MathWorks, Inc., Natick, MA). Copies of all custom code can be obtained from the authors.

## 3 Results

Figure 3 shows intelligibility scores for all conditions and experiments. Approximately equal overall intelligibility was achieved for intact and vocoded SiB due to our choice of SNRs for these conditions, based on extensive piloting. This allowed small differences in intelligibility to be normalized without loss of statistical power. Overall intelligibility was normalized to 60% for intact and vocoded SiB and to 90% for intact and vocoded SiQuiet, respectively (as described in Section 2.5), before examining the effects of vocoding on voicing, POA, and MOA confusion patterns.

**Figure 3.**
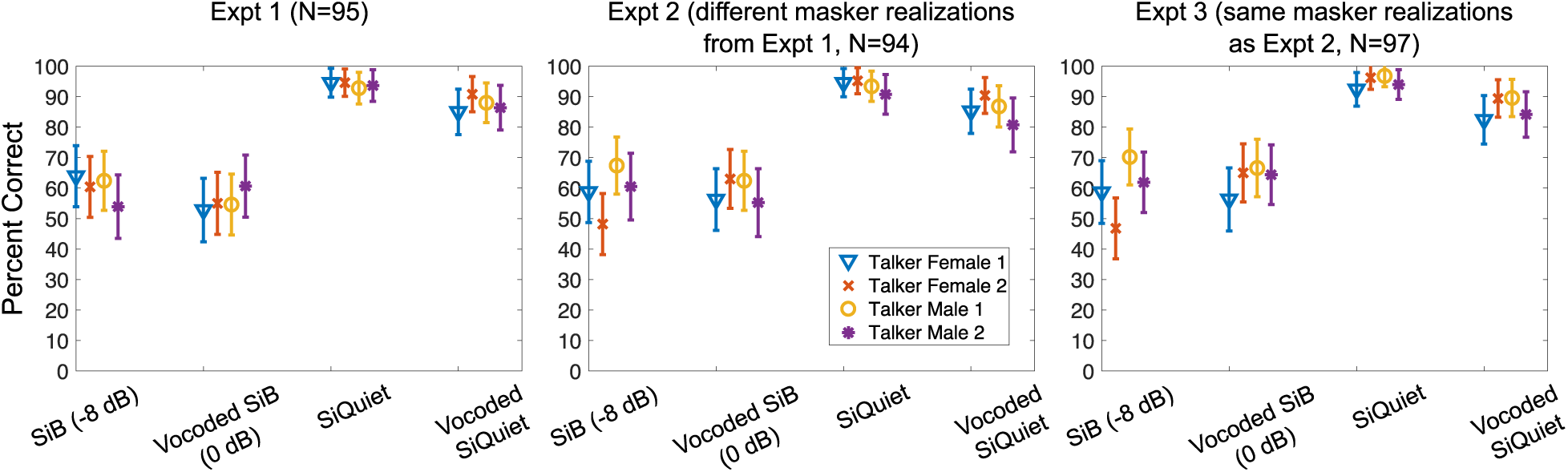
Overall intelligibility (mean and standard error) measured in the online consonant identification experiments for the different conditions and talkers. Approximately equal overall intelligibility was achieved across intact and vocoded SiB, and across intact and vocoded SiQuiet.

Given that our data were collected online, a few different data quality checks were conducted. The first of these examined whether subjects randomly chose a different consonant from what was presented when they made an error or if there was more structure in the data. As shown in Figure 4, percent errors in our data fall outside the distributions expected from random confusions. This result suggests that the error patterns in our data have a non-random structure, which supports the validity of our online-collected data. Moreover, there are small differences in the percent errors for voicing, place, and manner between intact and vocoded SiB and also between intact and vocoded SiQuiet. These differences were further investigated by quantifying full consonant confusion matrices for the voicing, place, and manner categories and examining the differences in these matrices across intact and vocoded conditions (Figs. 6, 7, and 8). This allowed us to obtain a richer characterization of the error patterns in consonant categorization (i.e., when an error was made, what consonant was reported instead of the consonant presented, and on what proportion of trials the alternative was reported) compared to the percent error scores shown in Figure 4.

**Figure 4.**
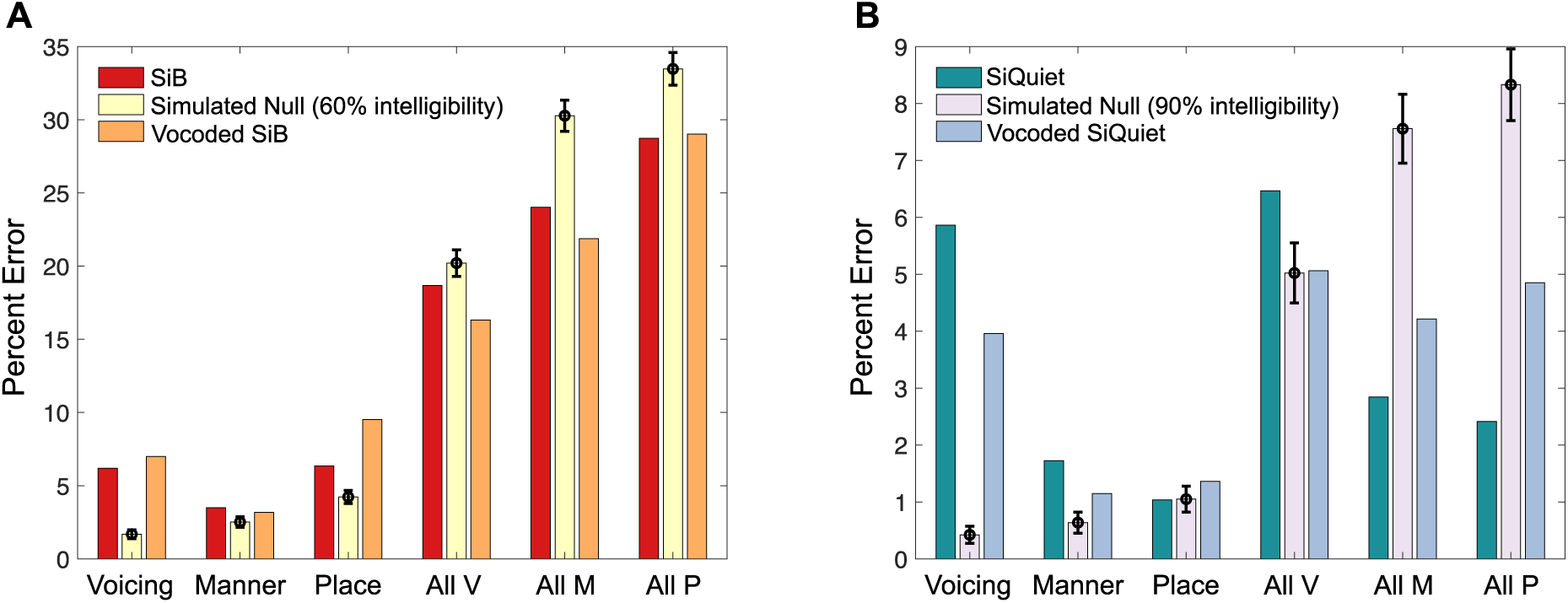
Percent errors (mean and standard deviation from experiment 1) for each phonetic category for intact and vocoded SiB (Panel A), and intact and vocoded SiQuiet (Panel B). The labels “Voicing”, “Manner”, and “Place” correspond to when the consonant reported differed from the consonant presented only in voicing, MOA, or POA, respectively. “All V”, “All M”, and “All P” correspond to when the consonant reported differed from the consonant presented in at least voicing, MOA, or POA, respectively [e.g., “All V” includes the following types of errors: (i) voicing only, (ii) voicing and MOA simultaneously, (iii) voicing and POA simultaneously, and (iv) voicing, MOA, and POA simultaneously]. The expected distribution of errors under the null hypothesis of random confusions was generated separately for Panels A and B, and with 1000 realizations each. Each realization of each null distribution was produced by generating a Bernoulli trial with “success” probability = 60% for Panel A, or 90% for Panel B, followed by uniform-random selection of a different consonant from what was presented if the trial outcome was “failure”.

To further test data quality, consonant confusions for the SiSSN condition were compared with previous lab-based findings since speech-shaped stationary noise is a commonly used masker in the phoneme confusion literature. Phatak and Allen (2007) found that for a given overall intelligibility, recognition scores varied across consonants. They identified three groups of consonants, “C1”, “C2”, and “C3” with low, high, and intermediate recognition scores, respectively in speech-shaped noise. Our online-collected data for SiSSN (Fig. 5A) closely replicate that key trend for the groups they identified after matching the SNR they used. Moreover, using a hierarchical clustering analysis (Ward Jr, 1963) of the consonant confusion matrix (pooled over samples) for SiSSN, perceptual “clusters” (i.e., sets where one consonant is confused most with another in the same set) were identified (shown as a dendrogram plot in Fig. 5B). The clusters identified here closely replicate the lab-based clustering results of Phatak and Allen (2007), further supporting the validity of our online data.

**Figure 5.**
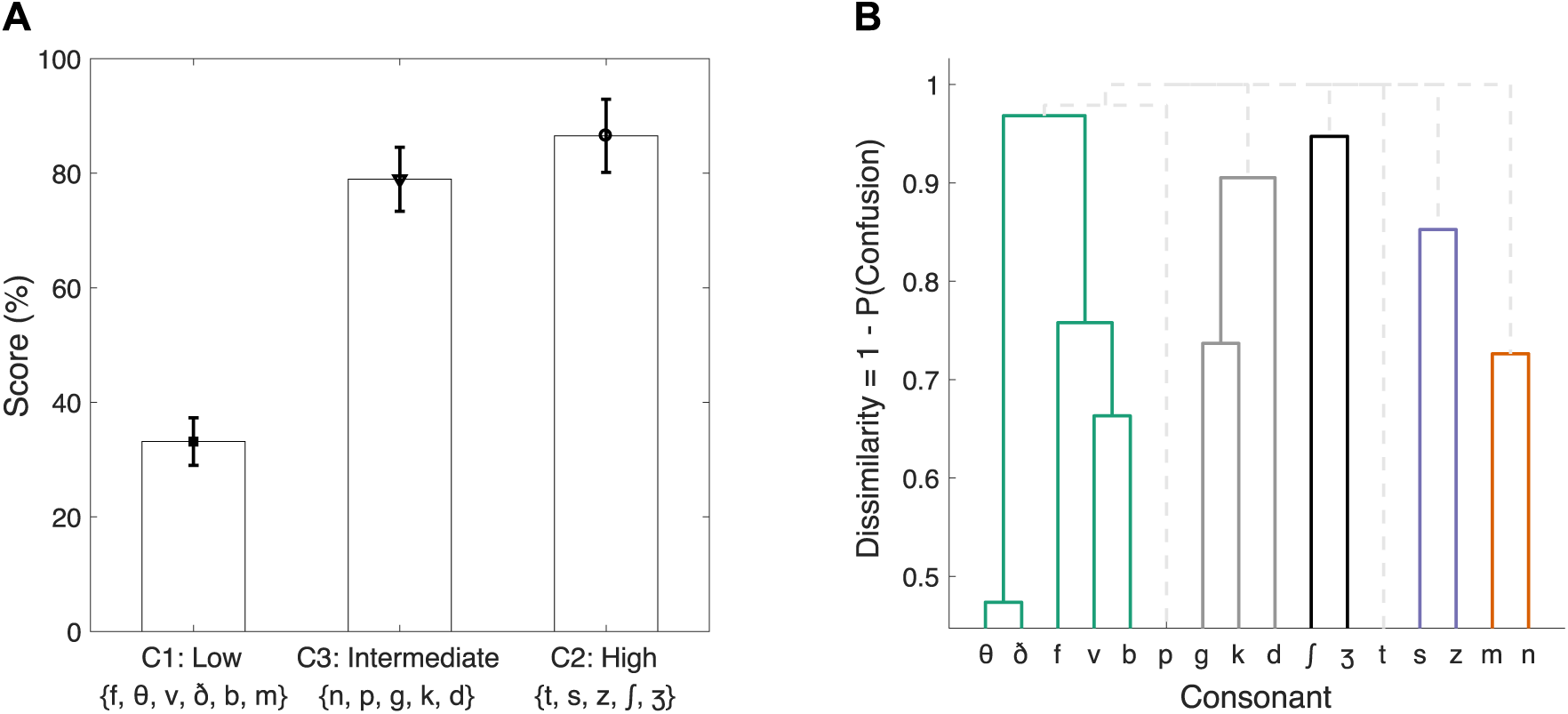
Consonant groups (Panel A) and confusion clusters (Panel B) for the speech in speech-shaped stationary noise (SiSSN) data from experiment 1. Panel A shows recognition scores for our SiSSN data for the three groups of consonants, “C1”, “C2”, and “C3” that Phatak and Allen (2007) identified as having low, high, and intermediate recognition scores, respectively, in speech-shaped noise (for a given overall intelligibility). Panel B shows the perceptual “clusters” (visualized as a dendrogram plot) identified with our SiSSN data. Each cluster is a set where one consonant is confused most with another in the same set. Clusters with greater than 3% probability of confusion share a color. For example, /θ/ and /ð/ form a cluster because they are more confused with each other than with the other consonants; moreover, while /θ/ and /ð/ are less confused with the cluster comprising /f/, /v/, and /b/ than with each other, they are even less confused with all the remaining consonants.

After verifying data quality, the hypothesis that confusion patterns would be the same for intelligibility-matched intact and envelope-vocoded speech in babble was tested. Figure 6 shows the results for voicing confusions. Vocoding altered the voicing percept for speech in babble by changing subject bias relative to an ideal observer. In particular, there was a greater tendency in the vocoded (versus intact) condition for the subject to be biased toward reporting an unvoiced consonant despite envelope and place cues being largely preserved. A detection-theoretic analysis (see Section 2.7) was used to quantify the decision boundary for the average subject’s perceptual decision on whether or not to reject the null hypothesis that an unvoiced consonant was presented. The bias or shift in this boundary relative to an unbiased ideal observer was then quantified and compared between intact and vocoded conditions. Intact-to-vocoded bias changes were found to be about 40%, 24%, and 19% in experiments 1, 2, and 3, respectively. Thus, the result that vocoding biases the voicing percept toward unvoiced consonants is replicated across experiments 1–3, supporting the idea that this bias effect is robust and generalizes across different babble instances. Note that the bias change between the intact and vocoded SiB conditions was observed even though the percent correct scores for the unvoiced and voiced categories were similar across these conditions (i.e., the diagonal entries in Fig. 6 are zero after statistical testing; for the precise number of errors, see Fig. 4). That is, while there were not a significantly different number of voicing errors after vocoding, the errors in the vocoded condition were biased toward reporting an unvoiced consonant even when a voiced consonant was presented. The errors in the intact condition were biased in the opposite direction, causing the total number of errors to be similar across the two conditions. This result suggests that the original TFS conveys important voicing information even when envelope cues are intact, since degrading the TFS led to a greater bias toward the percept of unvoiced consonants. This result also demonstrates that independent insight can be gained into the role of TFS cues from analyzing error patterns in consonant categorization rather than just examining transmission scores for the different phonetic categories.

**Figure 6.**
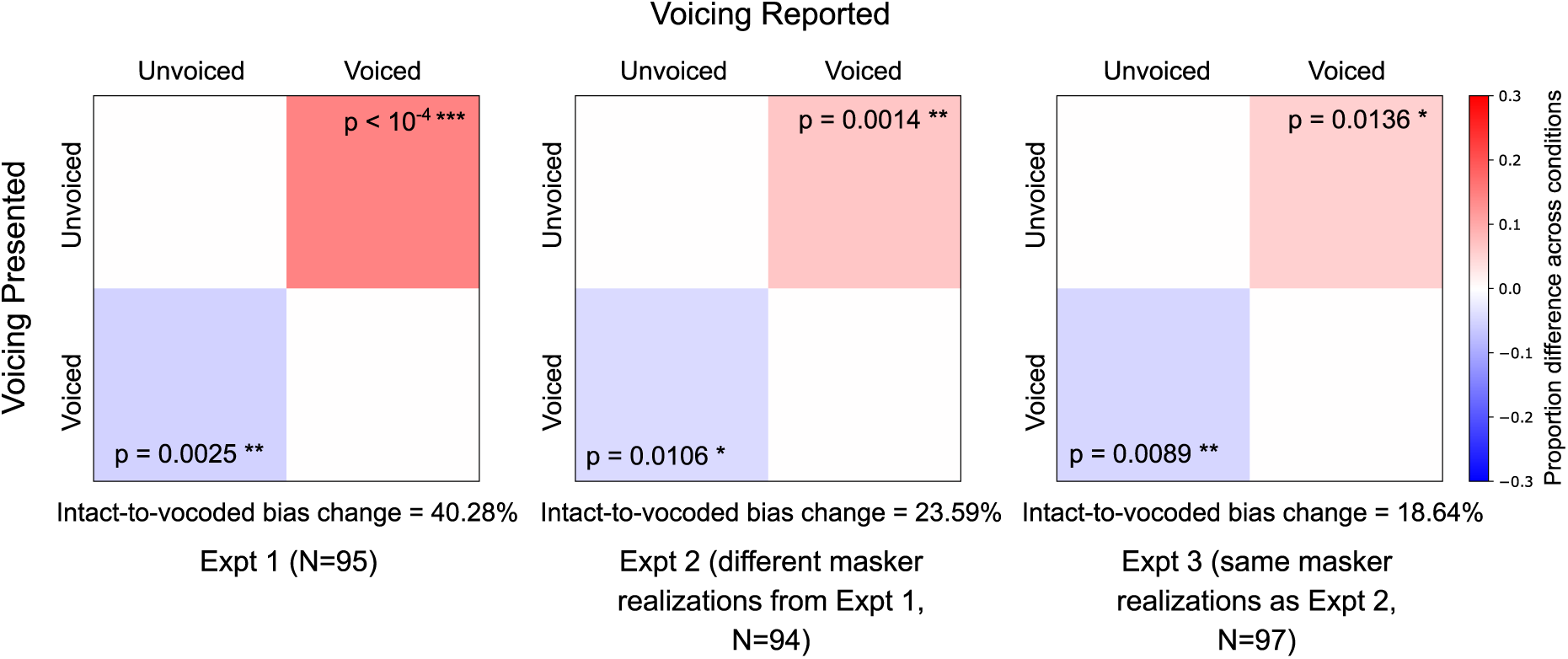
Voicing confusion-matrix differences (pooled over consonants and samples) between intact and vocoded SiB conditions (SiB - vocoded SiB). Overall intelligibility was matched at 60% before computing the differences across conditions. Only significant differences are shown, after permutation testing with multiple-comparison correction (5% FDR). Uncorrected p-values are also indicated for the individual matrix entries.

Figures 7 and 8 show the results from testing our hypothesis for POA and MOA confusions. Although significant differences were found in the POA and MOA confusion patterns between intact and vocoded SiB, the results were not consistent across experiments 1 and 2, which used different instances of babble (*R*^2^ = 2 × 10^*−*6^, *p* = 0.99 for POA, and *R*^2^ = 0.03, *p* = 0.44 for MOA). The results were replicated only when the stimuli were kept constant between experiments 2 and 3 (*R*^2^ = 0.85, *p* = 3.77 × 10^*−*13^ for POA, and *R*^2^ = 0.94, *p* = 1.44 × 10^*−*12^ for MOA). Note that the differences in POA and MOA confusions between intact and vocoded SiB could be due to either TFS or masker-instance effects; our goal behind using different masker instances across experiments 1 and 2 was to extract those effects that are not instance-specific but are rather due to a true effect of TFS. However, because the confusion-matrix differences for POA and MOA were not replicated across different masker instances, it is not possible to disambiguate between these two effects here. Nevertheless, the fact that these results did not generalize across different babble instances suggests that any effects of TFS on POA and MOA reception are weak when compared to differences across different samples of babble.

**Figure 7.**
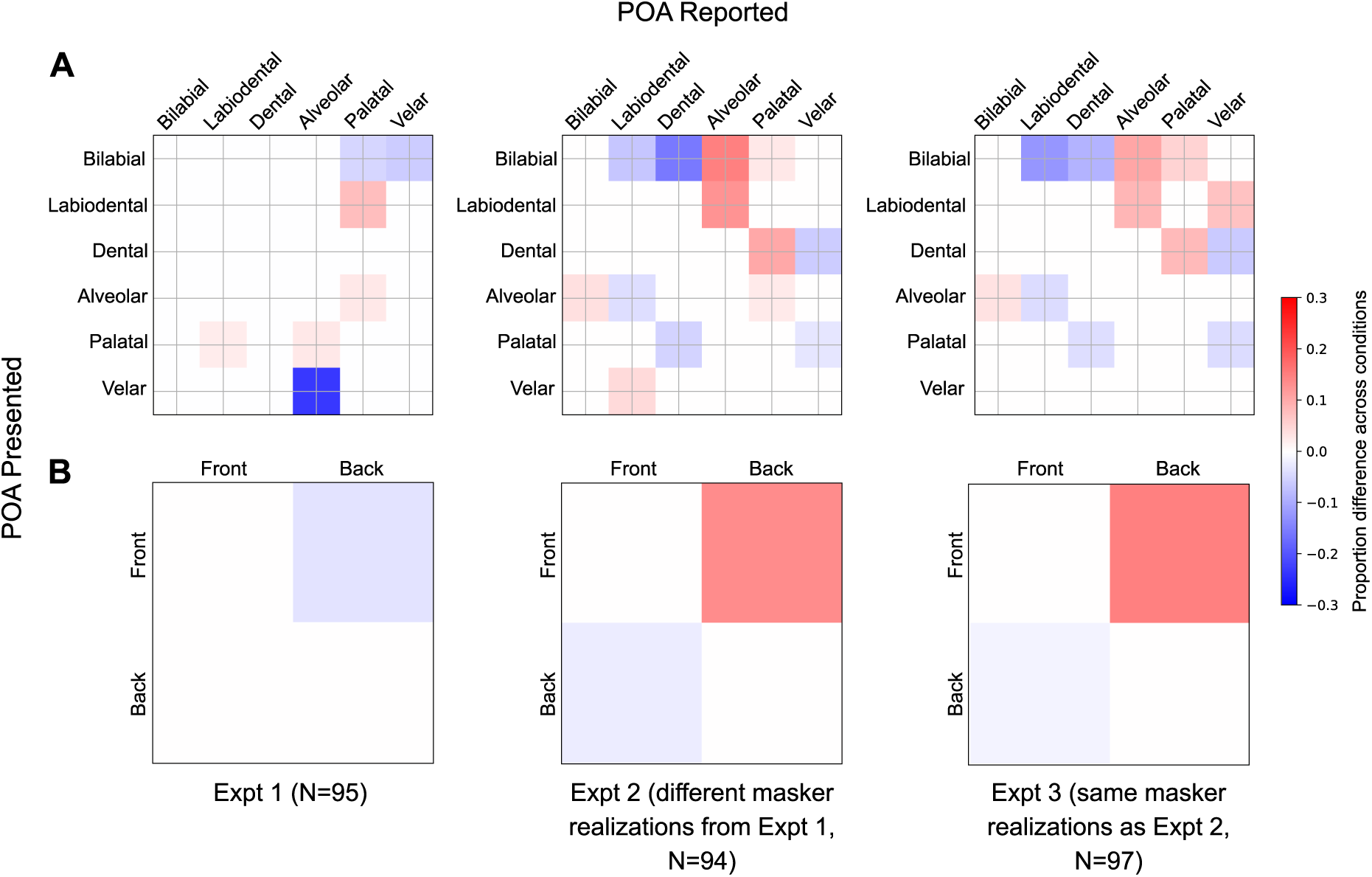
POA confusion-matrix differences (pooled over consonants and samples) between intact and vocoded SiB (SiB - vocoded SiB). Overall intelligibility was matched at 60% before computing the differences across conditions. Panel A shows full (5×5) matrices, whereas Panel B shows simplified (binary) matrices after collapsing over front versus back places of articulation. Only significant differences are shown, after permutation testing with multiple-comparison correction (5% FDR).

**Figure 8.**
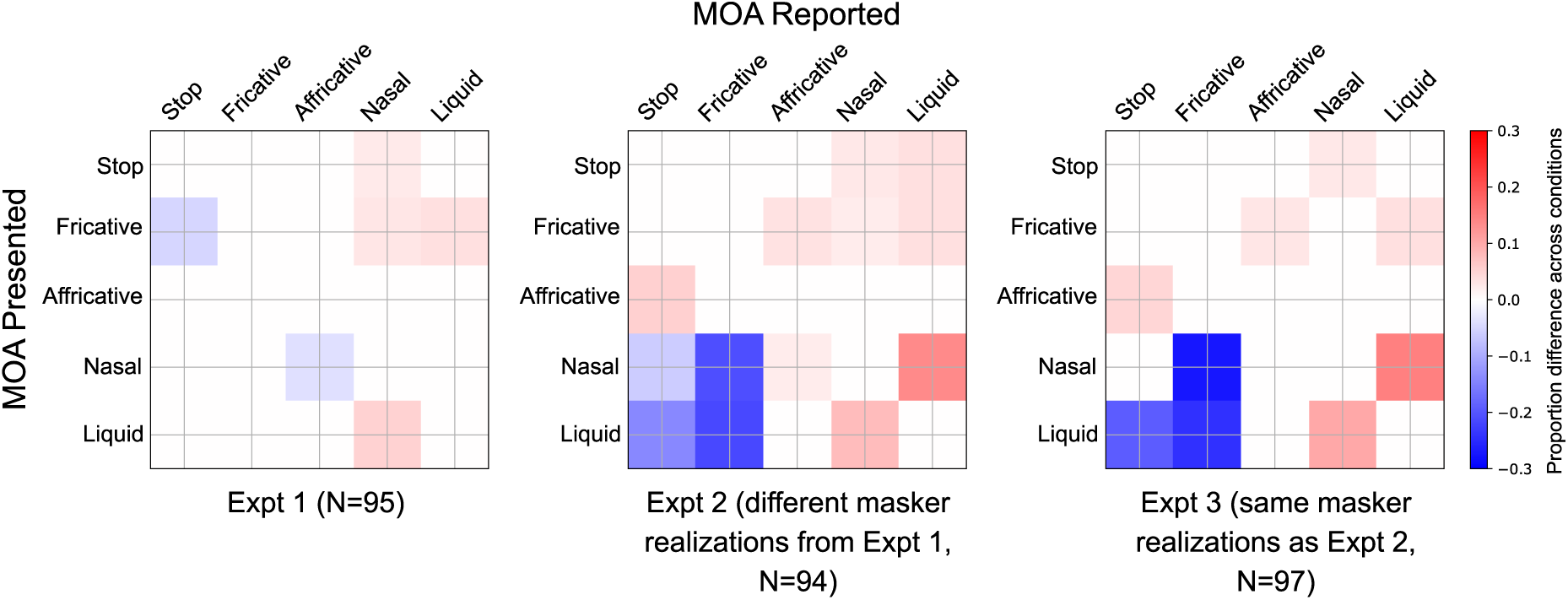
MOA confusion-matrix differences (pooled over consonants and samples) between intact and vocoded SiB (SiB - vocoded SiB). Overall intelligibility was matched at 60% before computing the differences across conditions. Only significant differences are shown, after permutation testing with multiple-comparison correction (5% FDR).

To test whether TFS conveys phonetic content beyond what is conveyed by envelopes for intact speech in quiet, the effect of vocoding on consonant confusion patterns for the SiQuiet condition was examined. This was done by computing confusion-matrix differences (after matching overall intelligibility at 90% and pooling across all experiments, consonants, and samples) between intact and vocoded SiQuiet for the voicing, POA, and MOA categories and performing permutation testing with multiple-comparison correction (5% FDR) to test for statistically significant differences (see Sections 2.5 and 2.6). The results indicate no significant effects of degrading TFS on either voicing, POA, or MOA confusions in quiet (figure not shown).

## 4 Discussion

The present study examined the influence of TFS on consonant confusion patterns by degrading TFS using high-resolution vocoding while controlling intelligibility to match that for intact stimuli. The results suggest that TFS is used to extract voicing content for intact speech in babble (i.e., even when redundant envelope cues are available). Moreover, this finding generalized across different babble instances. However, there were no significant vocoding effects on consonant confusions in quiet even after pooling data across all experiments; instead, overall intelligibility for vocoded SiQuiet was ∼90%.

The finding that TFS conveys voicing information beyond what is conveyed by envelopes for intact speech in babble is previously unreported to the best of our knowledge. This result deviates from the commonly held view that envelopes convey most speech content (Shannon et al., 1995). Several acoustic cues have been implicated in the categorization of consonant voicing, such as voice onset time (VOT), F0 at the onset of voicing, and the relative amplitude of any aspiration noise in the period between the burst release and the onset of voicing (Francis et al., 2008). Of these, VOT appears to be the dominant cue in quiet (Francis et al., 2008). However, listeners shift reliance to onset F0 when the VOT is ambiguous in the presence of noise (Winn et al., 2013; Holt et al., 2018). Our finding that vocoding alters the voicing percept in noise, but not quiet, is consistent with this result from the cue-weighting literature and can be attributed to impaired F0 cues resulting from TFS degradation in the vocoded (versus intact) SiB condition. Indeed, voiced sounds (unlike unvoiced) have quasi-periodic acoustic energy reflecting the quasi-periodic vibrations of the vocal folds; this periodicity has an F0 that is perceived as pitch (Rosen, 1992). Our finding that TFS is used to extract voicing content for intact speech in babble is consistent with the view that the pitch of complex sounds (with resolved harmonics) is coded either via TFS (Meddis and O’Mard, 1997; Moore et al., 2006) or via a combination of TFS and tonotopic place (Shamma and Klein, 2000; Oxenham et al., 2004). Indeed, psychophysical studies have found that melody perception (Moore and Rosen, 1979) and F0 discrimination (Houtsma and Smurzynski, 1990; Bernstein and Oxenham, 2006) are both better when conveyed by low-frequency resolved harmonics where the auditory nerve can robustly phase lock to the TFS (Johnson, 1980; Verschooten et al., 2015). Our results from directly manipulating TFS cues also corroborate previous correlational work relating model auditory-nerve TFS coding and voicing reception in noise (Swaminathan and Heinz, 2012). Other previous studies have suggested that low-frequency speech information is important for voicing transmission (Li and Loizou, 2008), but the experimental manipulations they used altered multiple cues including low-frequency place cues, slower envelopes, and possibly the masking of more basal regions by upward spread; this makes it difficult to unambiguously attribute their results to the role of TFS. In contrast, by isolating TFS manipulations in the present study, these limitations were overcome.

A potential contributor to the confusion-matrix differences between the intact and vocoded SiB conditions in the current study could be differences in the energetic masking of the target-speech envelope (due to differences in the acoustic SNR between these conditions). Future studies should explore this possibility further by testing whether there are significant changes in the pattern of consonant confusions for intact speech in babble as the input SNR is varied. However, there are several caveats to note when comparing confusion patterns across conditions with different intelligibility levels. It is possible that the data may exhibit floor or ceiling effects for a subset of the conditions. Furthermore, a measurement condition with high intelligibility will likely need a large number of trials and/or participants so that sufficient statistical power may be obtained to accurately estimate the pattern of confusions. Finally, the variance of the confusion-pattern estimates will differ between conditions with unmatched intelligibility; this should be accounted for during statistical analysis.

The current study found a strong babble-instance effect on POA and MOA confusion patterns. The effects of vocoding on these confusion patterns were not replicated when babble instances differed between experiments 1 and 2 but were replicated when instances were fixed across experiments 2 and 3. One explanation for the differences in confusion patterns across varying babble instances is that even though the average masker modulation spectrum was kept constant (the envelope of babble is dominated by low modulation frequencies; Viswanathan et al., 2021), there can be small variations in the modulation spectrum of the babble masker across instances within any given short time window. This, in turn, can cause variations in modulation masking across instances due to the relatively short duration of each consonant. Although not directly tested, hints of such effects of short-term envelope statistics were also found by Phatak and Grant (2012), where alterations of masker modulations produced less predictable effects on consonants than on vowels. In the present study, masker-instance effects on consonant perception were explicitly measured and confirmed. The role of short-term masker statistics should be further examined in future studies, perhaps using computational modeling to predict instance effects on consonant confusions from variations in modulation masking across short masker instances. Indeed, psychoacoustic literature on speech-in-noise perception (Bacon and Grantham, 1989; Stone and Moore, 2014), neurophysiological studies using EEG (Viswanathan et al., 2021), and the success of current speech intelligibility models (Dubbelboer and Houtgast, 2008; Relaño-Iborra et al., 2016) show that modulation masking (i.e., masking of the internal representation of temporal modulations in the target by distracting fluctuations from the background) is a key contributor to speech perception in noise.

The fact that no significant vocoding effects on consonant confusions in quiet were found, even after pooling data across experiments, is consistent with previous behavioral studies that suggested that speech content in quiet is mostly conveyed by envelopes (Shannon et al., 1995; Elliott and Theunissen, 2009) and with the success of envelope-based cochlear implants in quiet backgrounds (Wilson and Dorman, 2008). However, our finding that voicing cues are degraded in vocoded (versus intact) SiB has implications for current cochlear implants that do not appear to be able to provide usable TFS cues (Magnusson, 2011; Heng et al., 2011) because babble is a masker that is ubiquitous in everyday listening environments. Indeed, multi-talker babble, which has modulations spanning the range of modulations in the target speech, is a more ecological masker than either stationary noise (which has predominantly high-but not low-frequency modulations as are present in speech) or even narrowband syllabic-range AM modulations imposed on stationary noise (Viswanathan et al., 2021), as were used in previous studies (Gnansia et al., 2009; Swaminathan and Heinz, 2012; Winn et al., 2013; Holt et al., 2018). In addition to our finding here that TFS can convey important voicing cues, there is evidence from previous studies that TFS can aid in source segregation (Darwin, 1997; Oxenham and Simonson, 2009; Micheyl and Oxenham, 2010) and stronger representation of attended-speech envelopes in the brain (Viswanathan et al., 2021). The effect of TFS on segregation is reflected in the present study too, where the SNR for vocoded SiB had to be increased by 8 dB relative to intact SiB to match their respective overall intelligibility values. Taken together, these results suggest that patients with cochlear implants may benefit from improvements that allow these implants to provide usable TFS cues for speech recognition in everyday listening environments with multiple talkers or sound sources (Magnusson, 2011; Heng et al., 2011). This should be further examined in future studies using clinical populations.

One limitation of the current study is the use of isolated CV syllables (e.g., /ba/) rather than words commonly used in the English language (e.g., bat) to measure consonant categorization. However, the use of CV syllables allowed us to easily standardize the context across the different consonants (i.e., the vowel used was always /a/, and it always occurred after the consonant), thereby eliminating any confounds between the consonant used and condition effects (i.e., the effect of vocoding). One issue with standardizing context in this manner is that the effect of TFS may depend on the specific context used (i.e., C/a/). Thus, future work should explore whether such interaction effects exist. That being said, the C/a/ syllables were not presented in complete isolation; instead, a carrier phrase was used to help guide the listeners’ attention in a manner similar to natural running speech.

## 5 Conclusions

Using envelope-vocoding experiments that controlled for overall performance, the present study found evidence that TFS is used to extract voicing content for intact speech in babble (i.e., even when redundant envelope cues are available). This result was robustly replicated when babble instances were varied across independent experiments. Given that babble is a masker that is ubiquitous in everyday environments, this finding has implications for the design of assistive listening devices such as cochlear implants.

## 6 Acknowledgments

This research was supported by National Institutes of Health Grant Nos. F31DC017381 (to V.V.), R01DC009838 (to M.G.H.), and R01DC015988 (to B.G.S.-C.) and Office of Naval Research Grant No. N00014-20-12709 (to B.G.S.-C.). The authors would like to thank Hari Bharadwaj for access to online psychoacoustics infrastructure (Bharadwaj, 2021; Mok et al., 2021). We also thank Agudemu Borjigan, Andrew Sivaprakasam, François Deloche, Hari Bharadwaj, Ivy Schweinzger, Ravinderjit Singh, and Satyabrata Parida for valuable feedback on an earlier version of this manuscript. Finally, we thank Christian Lorenzi, Brian Moore, and an anonymous reviewer for their insightful and helpful comments.

## 7 Author Contributions

V.V., B.G.S.-C., and M.G.H. designed research; V.V. performed research; V.V. analyzed data; V.V. wrote the paper with edits from B.G.S.-C. and M.G.H.

## 8 Appendix

For completeness, the raw confusion matrices for all conditions and experiments are shown in Figure 9.

**Figure 9.**
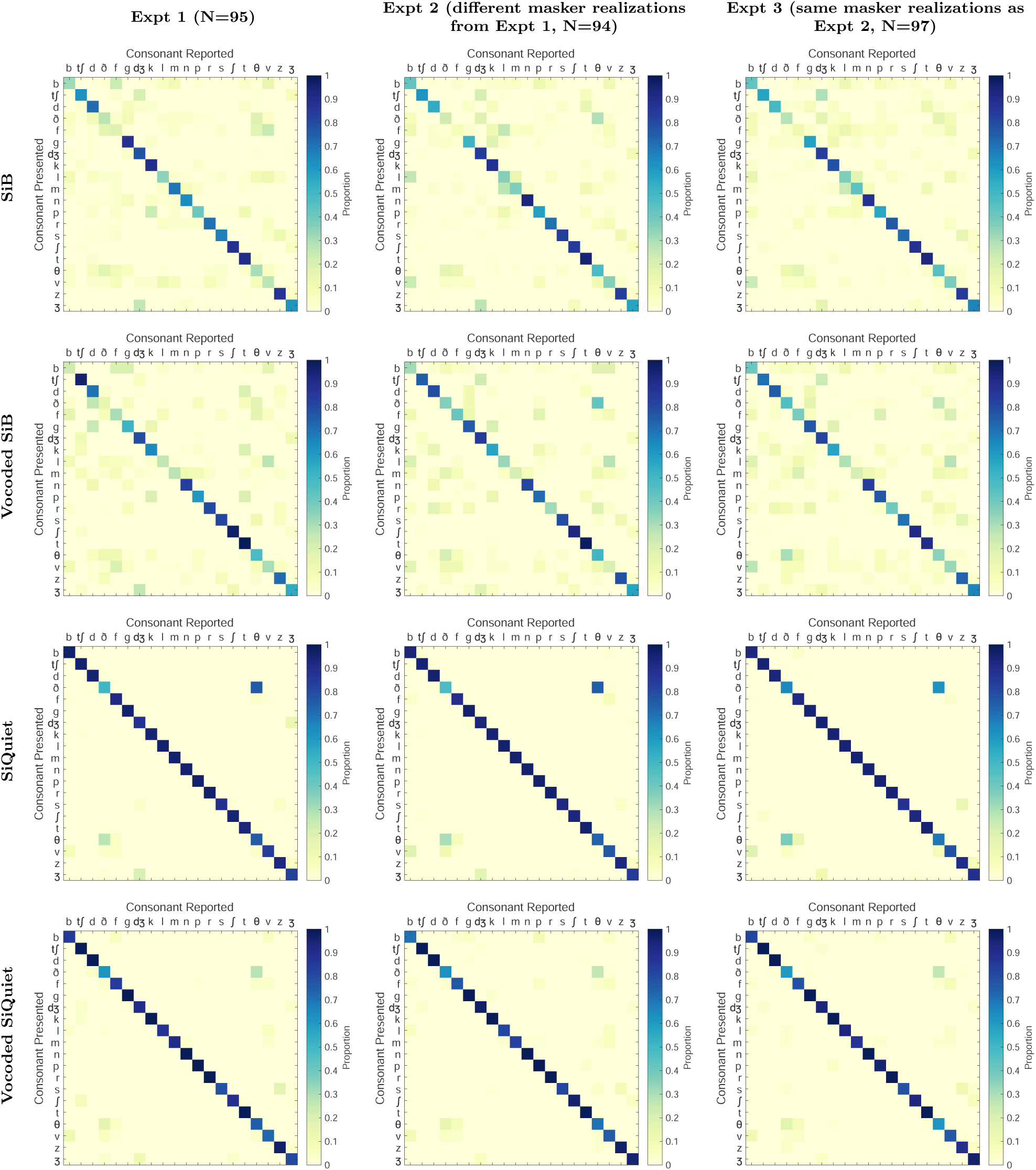
Raw confusion matrices for all conditions and experiments (pooled over samples). Overall intelligibility was 60% for the SiB and vocoded SiB conditions and 90% for the SiQuiet and vocoded SiQuiet conditions.

